## Supplementary_materials for "PointTree: Automatic and accurate reconstruction of long-range axonal projections of single-neuron"

Lin Cai *et al.*

**This PDF file includes:**

Figs. S1 to S7
Tables S1 to S2

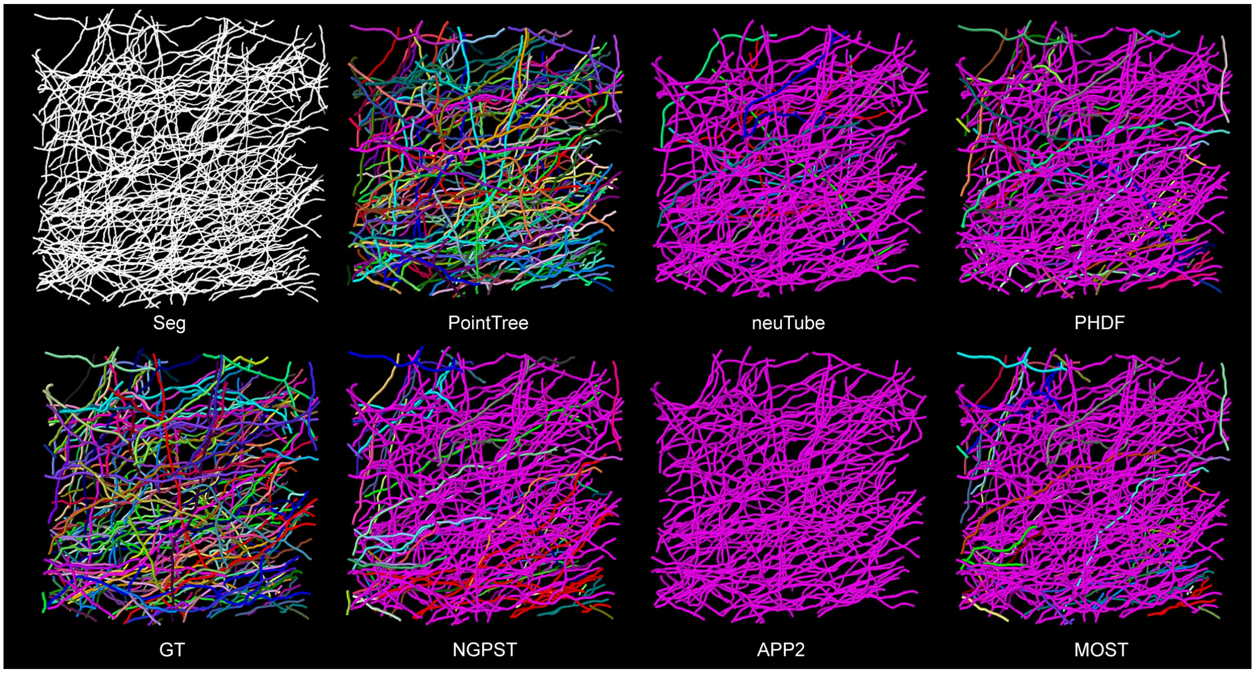


**Fig. S1. Comparison of PointTree and several skeleton-based methods for reconstructing the segmented image block derived from ground-truth skeletons**. The ground-truth skeletons are generated using GTree (a semi-automatic software) with manual modification. A series of Gaussian kernels with mean values equal to the coordinates of skeleton points are summed to obtain the corresponding probability image block. The segmented image block is finally generated using a threshold method.

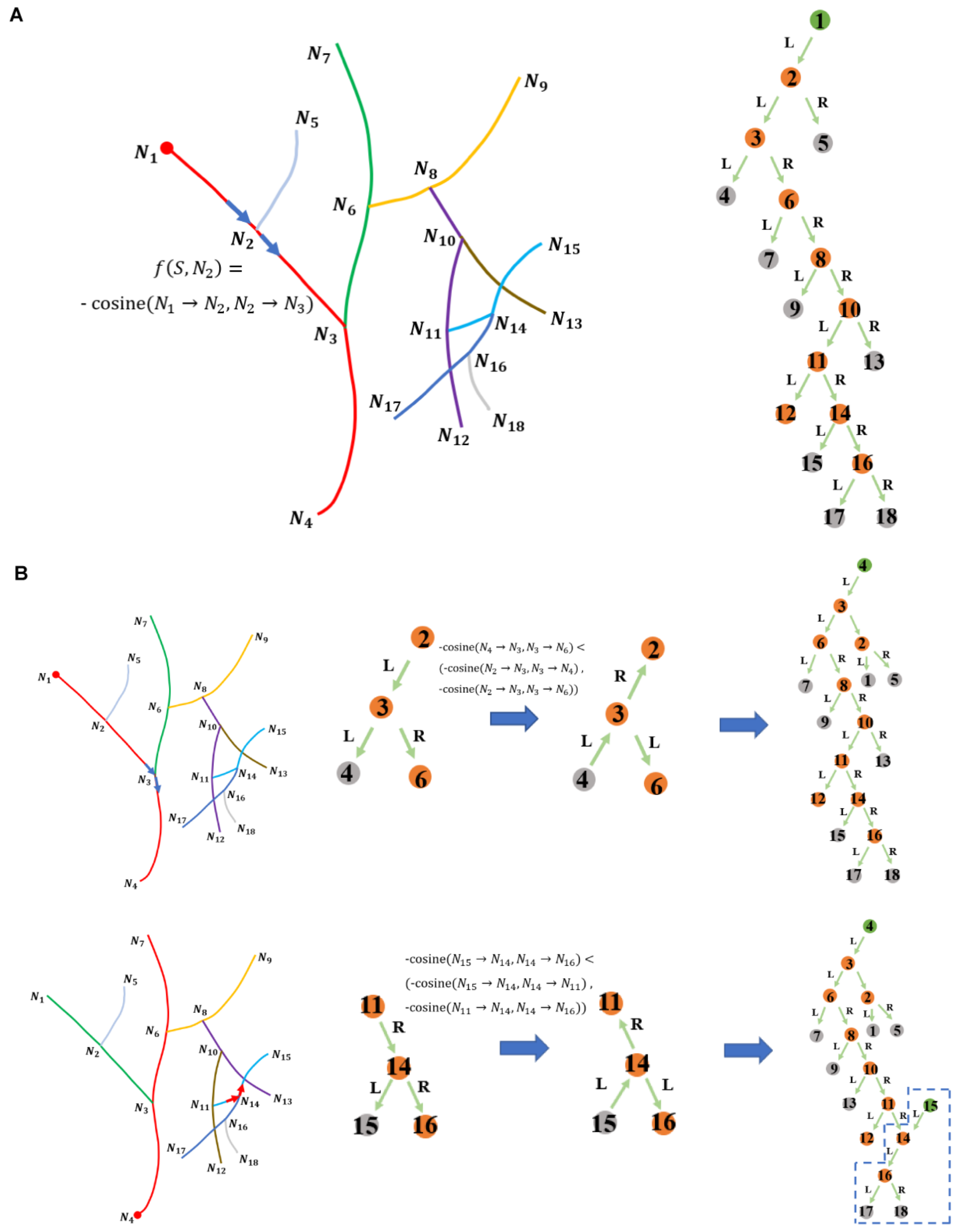


**Fig. S2. The generation of minimal information flow tree**. In (**A**), the calculation of the information flow score for a branch of neurites with root node 1 is illustrated. The reconstructed skeletons are transformed into a binary structure based on the root node, and the angles with respect to branching nodes labeled with brown circles determine the information flow. These angles will change when the root node changes. The angle of a branching node is formed by its father and child nodes, as exemplified by N2 node. In (**B**), the optimization of tree structure to minimize the total information flow score is demonstrated. It shows that decreasing the information flow leads to a more proper tree structure. The second row of (**B**) provides an example of decomposing tree structure into two individual parts.

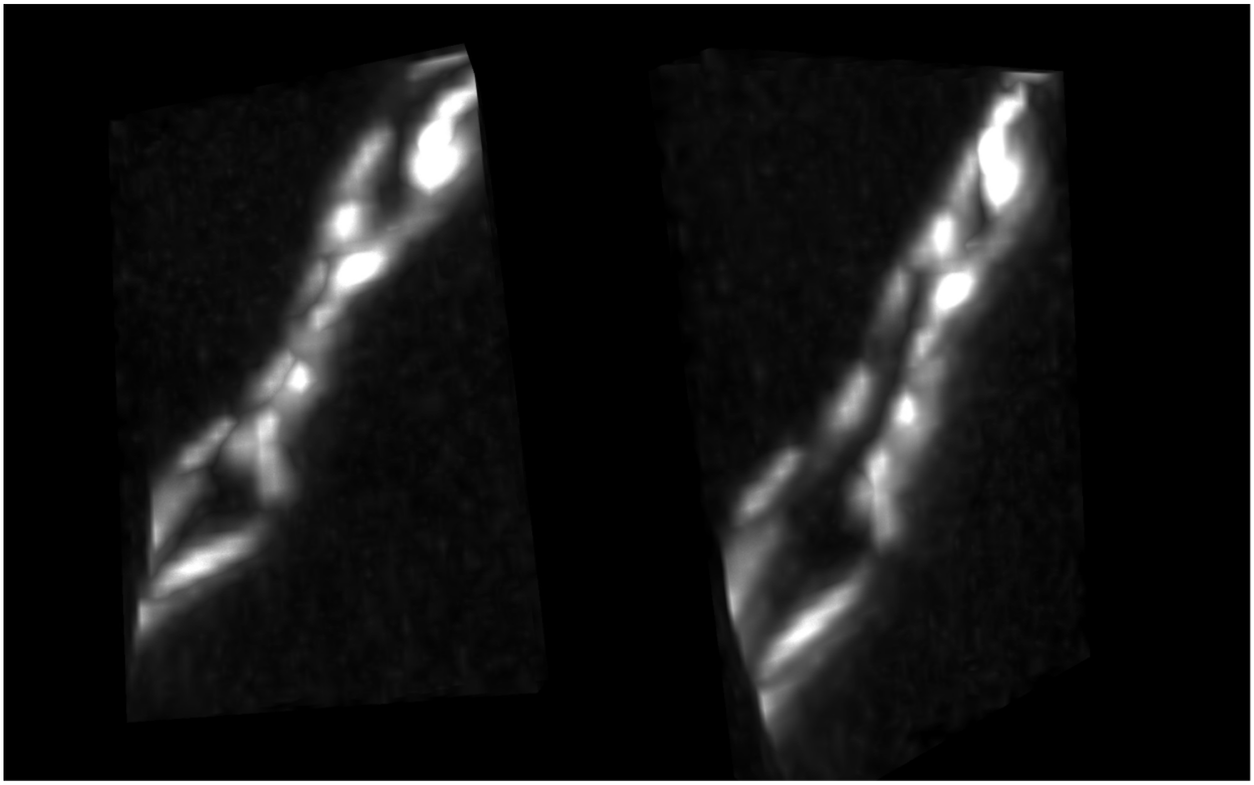

**Fig. S3. Visualization of two parallel neurites with different viewing angles**. When the two parallel neurites are in close proximity, they can be distinguished by visualizing them from different angles.

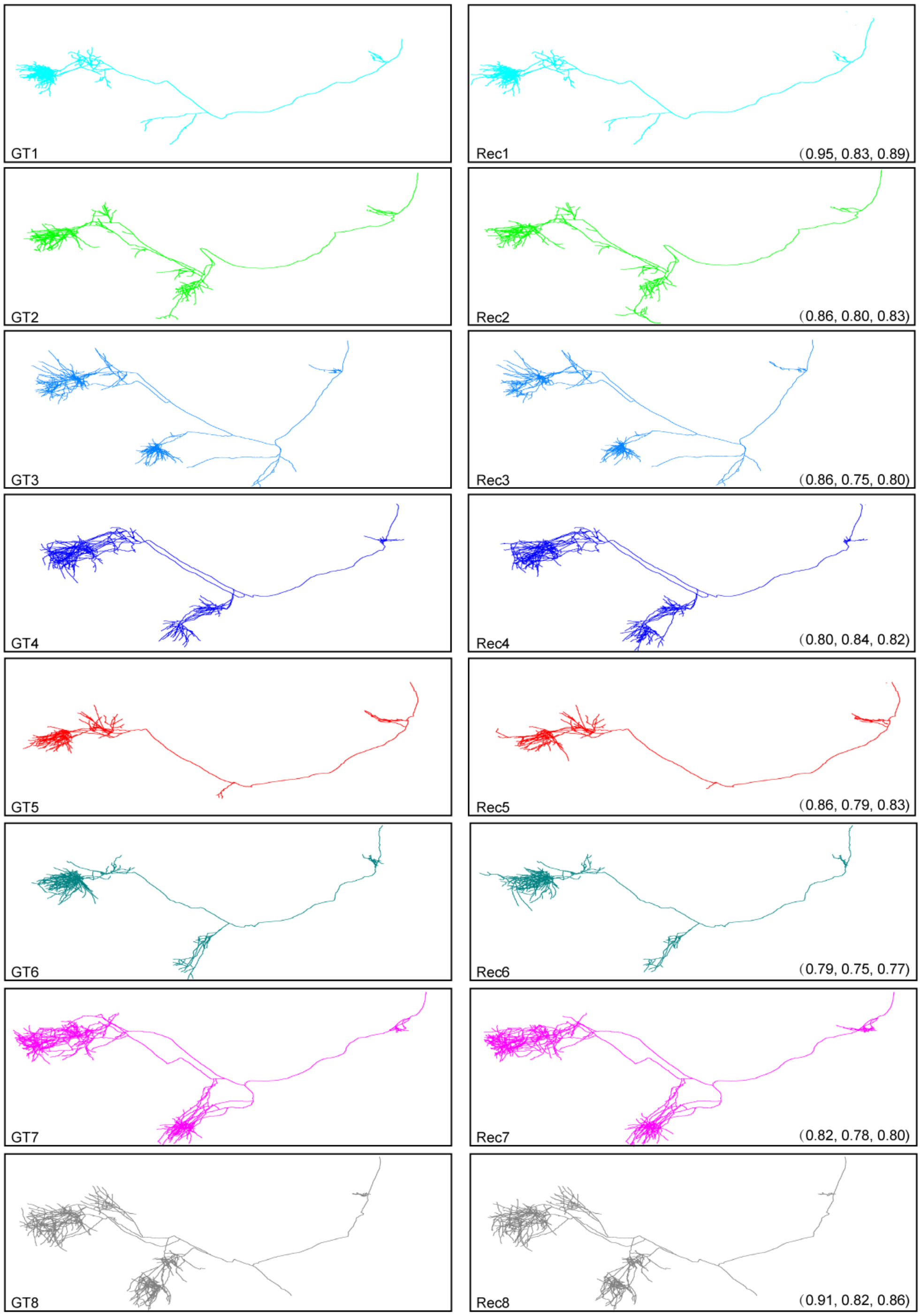


**Fig. S4**. **Reconstruction of long-range axonal projections.** The reconstructions were performed using semi-automatic methods with manual modification (GTree, left column) and automatic methods (PointTree, right column). The semi-automatic reconstruction is considered the ground-truth reconstruction for quantifying the accuracy of the automatic reconstruction. In the right column, each panel includes a set of quantitative evaluation indexes in the bottom-right corner, which consist of precision, recall, and f1-score.

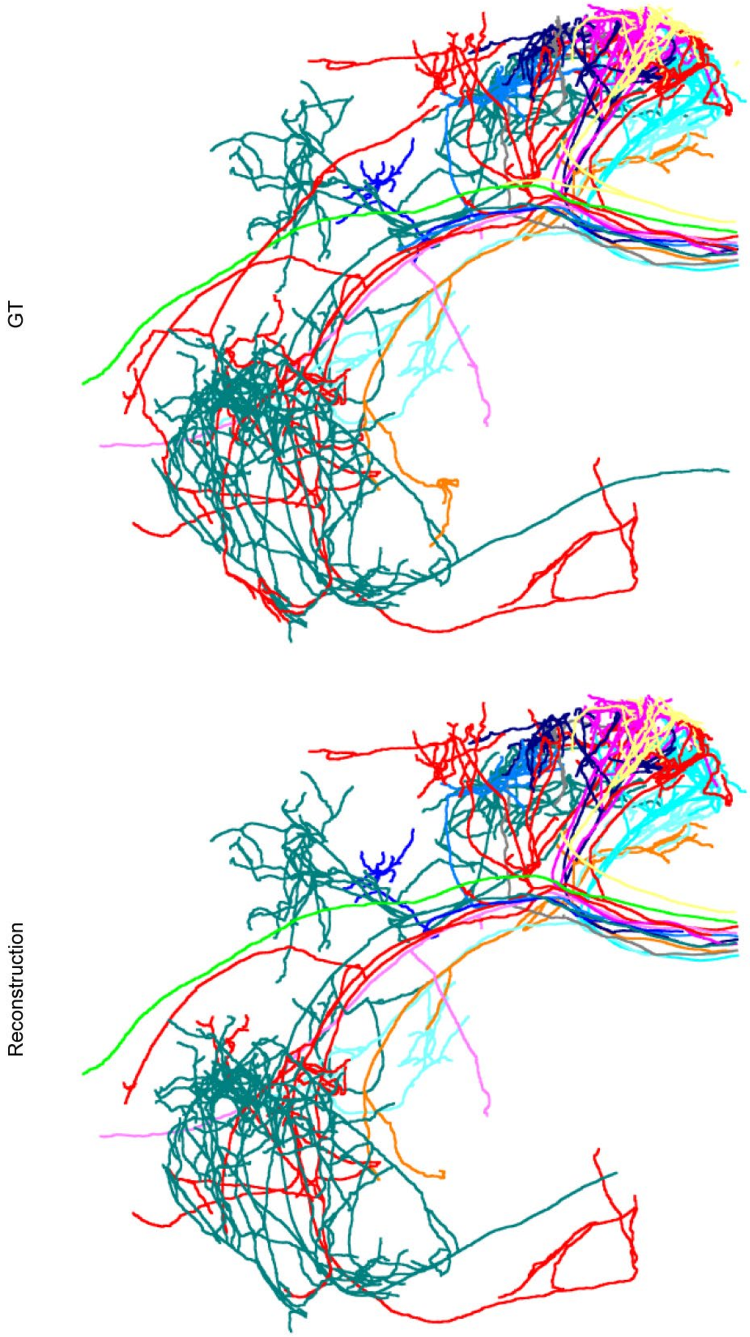


**Fig. S5**. **Reconstruction from different dataset.** Axonal reconstructions were generated from the image blocks (10739×11226×3921) collected using the Limo system. The upper portion represents the ground-truth reconstruction, which includes data from 13 neurons. The automatic reconstruction (shown at the bottom) closely matches the ground-truth reconstruction. A quantitative evaluation of the automatic reconstruction is presented in Table S1.

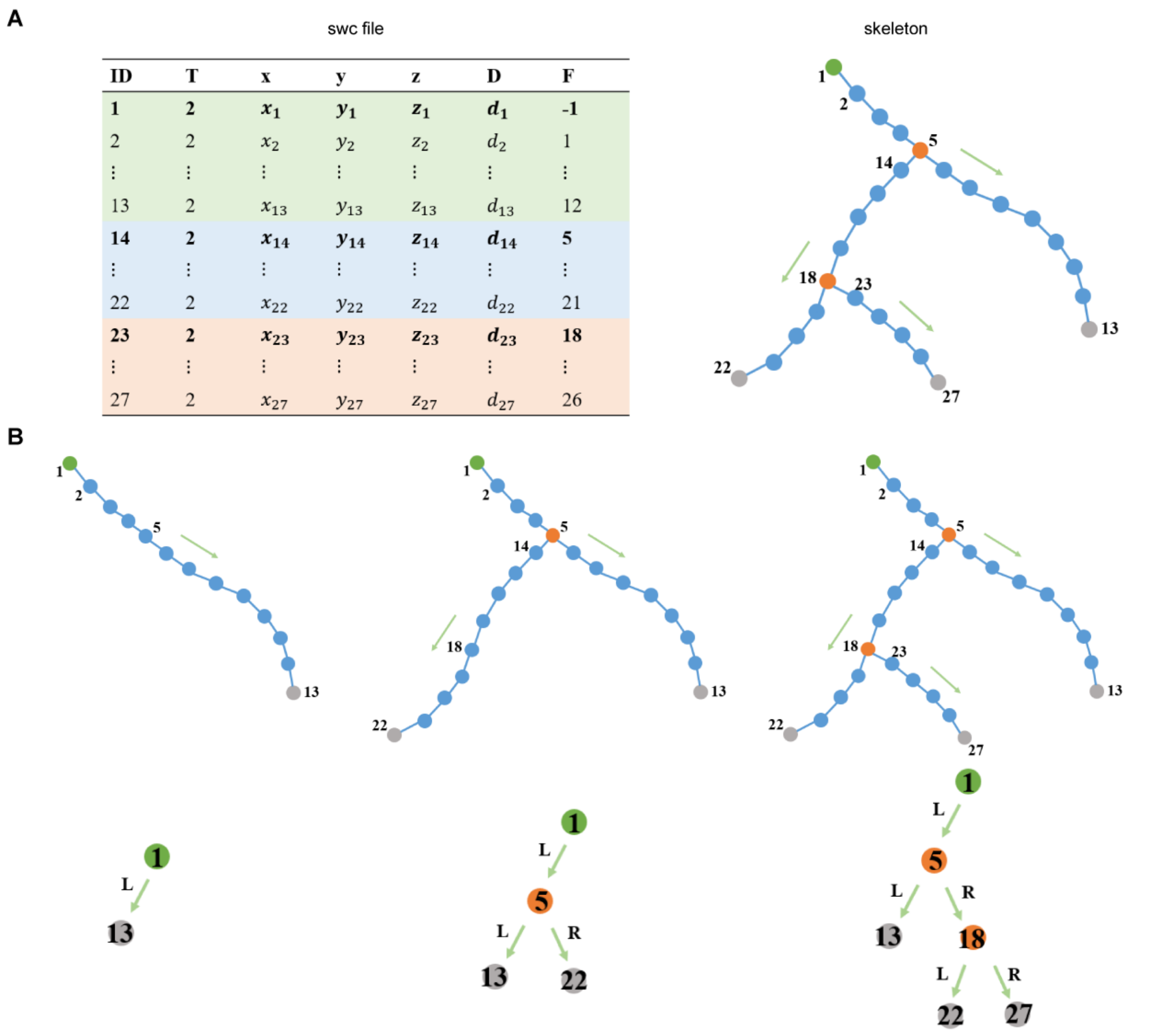


| ID | T | x | y | z | D | F |
|---|---|---|---|---|---|---|
| **1** | **2** | $\boldsymbol{x_1}$ | $\boldsymbol{y_1}$ | $\boldsymbol{z_1}$ | $\boldsymbol{d_1}$ | **-1** |
| 2 | 2 | $x_2$ | $y_2$ | $z_2$ | $d_2$ | 1 |
| ⋮ | ⋮ | ⋮ | ⋮ | ⋮ | ⋮ | ⋮ |
| 13 | 2 | $x_{13}$ | $y_{13}$ | $z_{13}$ | $d_{13}$ | 12 |
| **14** | **2** | $\boldsymbol{x_{14}}$ | $\boldsymbol{y_{14}}$ | $\boldsymbol{z_{14}}$ | $\boldsymbol{d_{14}}$ | **5** |
| ⋮ | ⋮ | ⋮ | ⋮ | ⋮ | ⋮ | ⋮ |
| 22 | 2 | $x_{22}$ | $y_{22}$ | $z_{22}$ | $d_{22}$ | 21 |
| **23** | **2** | $\boldsymbol{x_{23}}$ | $\boldsymbol{y_{23}}$ | $\boldsymbol{z_{23}}$ | $\boldsymbol{d_{23}}$ | **18** |
| ⋮ | ⋮ | ⋮ | ⋮ | ⋮ | ⋮ | ⋮ |
| 27 | 2 | $x_{27}$ | $y_{27}$ | $z_{27}$ | $d_{27}$ | 26 |

**Fig. S6**. **Tree structure derived from SWC file that records the axonal reconstruction.** (**A**) shows an input swc file and its corresponding skeleton structure. (**B**) shows how the reconstructed skeletons are converted to a binary tree structure.

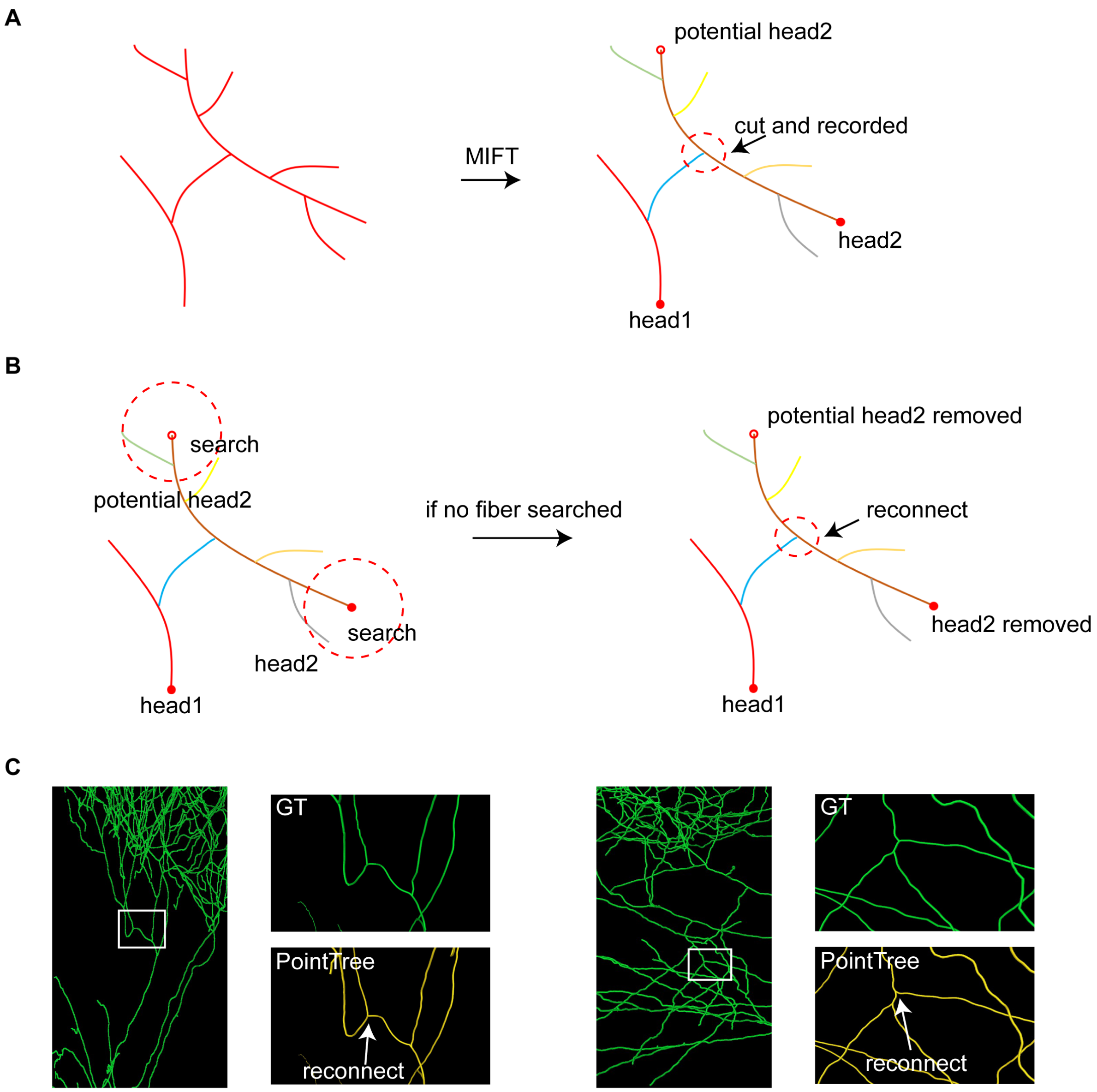


**Fig. S7. Post-processing for structures violating the MIFT rule.** (**A**) MIFT criterion will incorrectly split neurites with sharp directional changes into two branches, but the splitting location is explicitly recorded during this process. (**B**) Our algorithm searches for connectable neurites around the head nodes identified by MIFT. If no connectable neurites are found for both head nodes, the algorithm will reconnect them based on the recorded splitting points to prevent isolated neurite fragments. (**C**) presents two real examples violating the MIFT criterion. Through post-processing, PointTree successfully reconnects the split branches back to the correct neurites.

**Table S1.** Quantitative metrics comparing ground-truth and reconstructed neurons which are presented in Supplementary Fig.S6.

| ID | Precision | Recall | F1-Score |
|---|---|---|---|
| 1 | 1.00 | 0.92 | 0.95 |
| 2 | 1.00 | 1.00 | 1.00 |
| 3 | 0.98 | 0.76 | 0.86 |
| 4 | 1.00 | 0.82 | 0.90 |
| 5 | 1.00 | 0.77 | 0.87 |
| 6 | 1.00 | 0.92 | 0.96 |
| 7 | 0.96 | 0.75 | 0.84 |
| 8 | 1.00 | 0.87 | 0.93 |
| 9 | 1.00 | 0.82 | 0.90 |
| 10 | 1.00 | 0.96 | 0.98 |
| 11 | 1.00 | 0.99 | 0.99 |
| 12 | 1.00 | 0.77 | 0.87 |
| 13 | 1.00 | 0.90 | 0.95 |
| 14 | 0.99 | 0.87 | 0.93 |

**Table S2.** Time cost of three modules in the entire reconstruction for two testing datasets shown in Figure 5 and Fig.S6.

| block number (size: 512×512×512) | Points clustering (mins) | Clusters connection (mins) | Reconstruction merging (mins) |
|---|---|---|---|
| 254 | 23 | 18 | 3 |
| 821 | 22 | 35 | 3 |
